# QSMRim-Net: Imbalance-Aware Learning for Identification of Chronic Active Multiple Sclerosis Lesions on Quantitative Susceptibility Maps

**DOI:** 10.1101/2022.01.31.478482

**Authors:** Hang Zhang, Thanh D. Nguyen, Jinwei Zhang, Melanie Marcille, Pascal Spincemaille, Yi Wang, Susan A. Gauthier, Elizabeth M. Sweeney

## Abstract

**Background and Purpose:** Chronic active multiple sclerosis (MS) lesions are characterized by a paramagnetic rim at the edge of the lesion and are associated with increased disability in patients. Quantitative susceptibility mapping (QSM) is an MRI technique that is sensitive to chronic active lesions, termed rim+ lesions on the QSM. We present QSMRim-Net, a data imbalance-aware deep neural network that fuses lesion-level radiomic and convolutional image features for automated identification of rim+ lesions on QSM.

**Methods:** QSM and T2-weighted-Fluid-Attenuated Inversion Recovery (T2-FLAIR) MRI of the brain were collected at 3T for 172 MS patients. Rim+ lesions were manually annotated by two human experts, followed by consensus from a third expert, for a total of 177 rim+ and 3986 rim negative (rim-) lesions. Our automated rim+ detection algorithm, QSMRim-Net, consists of a two-branch feature extraction network and a synthetic minority oversampling network to classify rim+ lesions. The first network branch is for image feature extraction from the QSM and T2-FLAIR, and the second network branch is a fully connected network for QSM lesion-level radiomic feature extraction. The oversampling network is designed to increase classification performance with imbalanced data.

**Results:** On a lesion-level, in a five-fold cross validation framework, the proposed QSMRim-Net detected rim+ lesions with a partial area under the receiver operating characteristic curve (pROC AUC) of 0.760, where clinically relevant false positive rates of less than 0.1 were considered. The method attained an area under the precision recall curve (PR AUC) of 0.704. QSMRim-Net out-performed other state-of-the-art methods applied to the QSM on both pROC AUC and PR AUC. On a subject-level, comparing the predicted rim+ lesion count and the human expert annotated count, QSMRim-Net achieved the lowest mean square error of 0.98 and the highest correlation of 0.89 (95% CI: 0.86, 0.92).

**Conclusion:** This study develops a novel automated deep neural network for rim+ MS lesion identification using T2-FLAIR and QSM images.

## 1. INTRODUCTION

Multiple sclerosis (MS) is an inflammatory disease of the central nervous system, characterized by lesions in the brain and spinal cord [1]. A particular type of multiple sclerosis (MS) lesion, called a chronic active lesion, is characterized by an iron-enriched rim of activated macrophages and microglia in histopathology studies [2–5]. Chronic active lesions are visible with in-vivo susceptibility magnetic resonance imaging (MRI) techniques, where these lesions show a paramagnetic rim [2, 4, 6–15] on the edge. The presence of chronic active multiple sclerosis lesions is associated with a more severe disease course [14, 16–19] and there is currently much interest in using these lesions as an imaging biomarker.

In studies of chronic active MS lesions on MRI, lesions are typically identified on the T2-weighted-Fluid-Attenuated Inversion Recovery (T2-FLAIR) image and then are determined to be chronic active through visual inspection on susceptibility imaging. This process is time consuming and prone to inter- and intra-rater variability [20, 21]. For these lesions to be further studied at a large scale and translated into clinical practice, there is a great need for automated methods to identify chronic active MS lesions.

Quantitative susceptibility mapping (QSM) is an MRI technique that provides in vivo quantification of magnetic susceptibility changes related to iron deposition [22–24]. QSM identifies chronic active MS lesions as lesions with a hyperintense rim [25–27], which are termed QSM rim positive (rim+) lesions. We propose an automated method to identify QSM rim+ lesions, QSMRim-Net, using QSM and T2-FLAIR images of the brain. Our method is a deep convolutional neural network which consists of a two-branch network that fuses QSM and T2-FLAIR imaging features derived from a deep residual network [28] with lesion-level radiomic features from the QSM [29], In addition, a Synthetic Minority Oversampling TEchnique (SMOTE)-based oversampling network (DeepSMOTE) is developed to alleviate data imbalance issue caused by the small number of rim+ lesions. This is the first method proposed in the literature to identify rim+ lesions using QSM and the first method to fuse convolutional imaging features with radiomic features. Furthermore, QSMRim-Net with DeepSMOTE is the first end-to-end deep neural network that can be trained with online minority oversampling for rim+ lesion classification.

Two previous methods have been developed to identify chronic active MS lesions on phase imaging [30, 31]. RimNet [30] uses convolutional features, while APRL [49] uses radiomic features. To put the performance of QSMRim-Net into context, we compare it to these two methods applied to the QSM using both lesion-level and patient-level performance metrics.

## 2. MATERIALS AND METHODS

### 2.1 Dataset

#### 2.1.1 MRI image acquisition and preprocessing

QSMRim-Net was evaluated on an MS imaging dataset collected at Weill Cornell (Table 1). The dataset consists of 172 MS patients enrolled in an ongoing prospective database for MS research. The database was approved by the local Institutional Review Board and written informed consent was obtained from all patients prior to their entry into the database.

The imaging was performed on a 3T Magnetom Skyra scanner (Siemens Medical Solutions, Malvern, PA, USA). The Siemens scanning protocol consisted of the following sequences: 1) 3D sagittal T1-weighted (T1w) MPRAGE: Repetition Time (TR)/Echo Time (TE)/Inversion Time (TI) = 2300/2.3/900 ms, flip angle (FA) = 8°, GRAPPA parallel imaging factor (R) = 2, voxel size = 1.0 × 1.0 × 1.0 mm3; 2) 2D axial T2-weighted (T2w) turbo spin echo: TR/TE = 5840/93 ms, FA = 90°, turbo factor = 18, R = 2, number of signal averages (NSA) = 2, voxel size = 0.5 × 0.5 × 3 mm3; 3) 3D sagittal fat-saturated T2w fluid attenuated inversion recovery (T2-FLAIR) SPACE: TR/TE/TI = 8500/391/2500 ms, FA = 90°, turbo factor = 278, R = 4, voxel size = 1.0 × 1.0 × 1.0 mm3. For axial 3D multi-echo GRE sequence for QSM: axial field of view (FOV) = 24 cm, TR/TE1/ΔTE = 48.0/6.3/4.1 ms, number of TEs = 10, FA = 15°, R = 2, voxel size = 0.75 x 0.93 x 3 mm3, scan time = 4.2 minutes. QSM images were reconstructed by MEDI-0 [32] algorithm from multi-echo GRE data. T2-FLAIR images were then preprocessed using the FSL toolbox [33]. We applied N4 inhomogeneity correction algorithm to the acquired images and linearly co-registered T2-FLAIR images to the magnitude space of QSM.

#### 2.1.2 Lesion segmentation and rim+ lesion annotation

T2-FLAIR lesion masks were created for all patients in the dataset. These masks were obtained by segmenting the T2-FLAIR image using the LST-LPA algorithm in the LST toolbox version 3.0.0 (www.statisticalmodelling.de/lst.html) [34], followed by manual editing, and finalized by the consensus of two expert raters. Confluent lesions may occur when pathologically distinct lesions grow close to each other and form a large spatially connected lesion. These confluent lesions in the dataset were identified, then broken up and labeled by a human expert. After lesion segmentation and confluent lesion separation, a total of 4,163 individual lesions were identified. Masks were further edited on the QSM image to ensure that these masks matched the lesion on QSM. An overview of the annotation process is shown in Fig. 2.

**Fig. 1:**
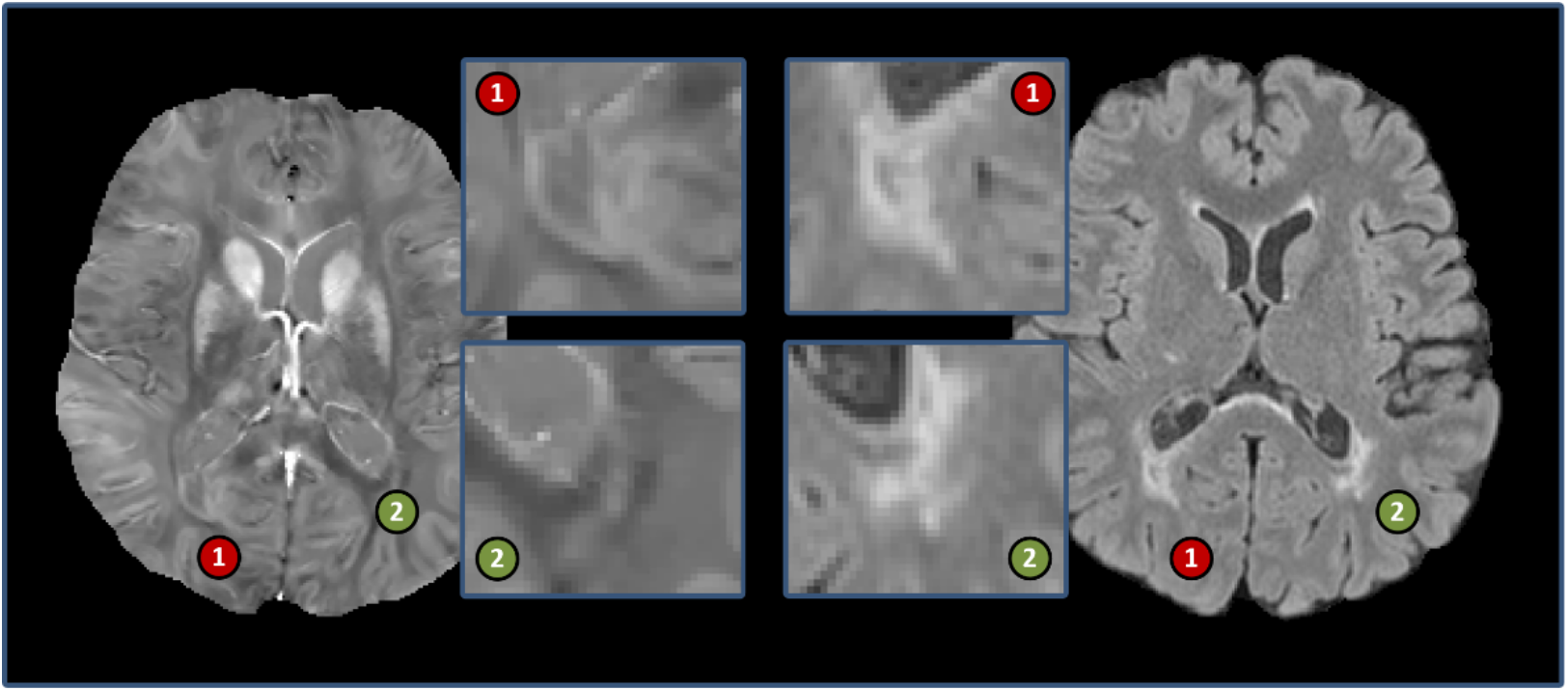
Example of MS lesions on an axial slice of the QSM (left) and corresponding axial slice of the T2-FLAIR (right). The digit 1 marked with red indicates a rim+ lesion and the digit 2 marked with green indicates a rim-lesion.

**Fig. 2:**
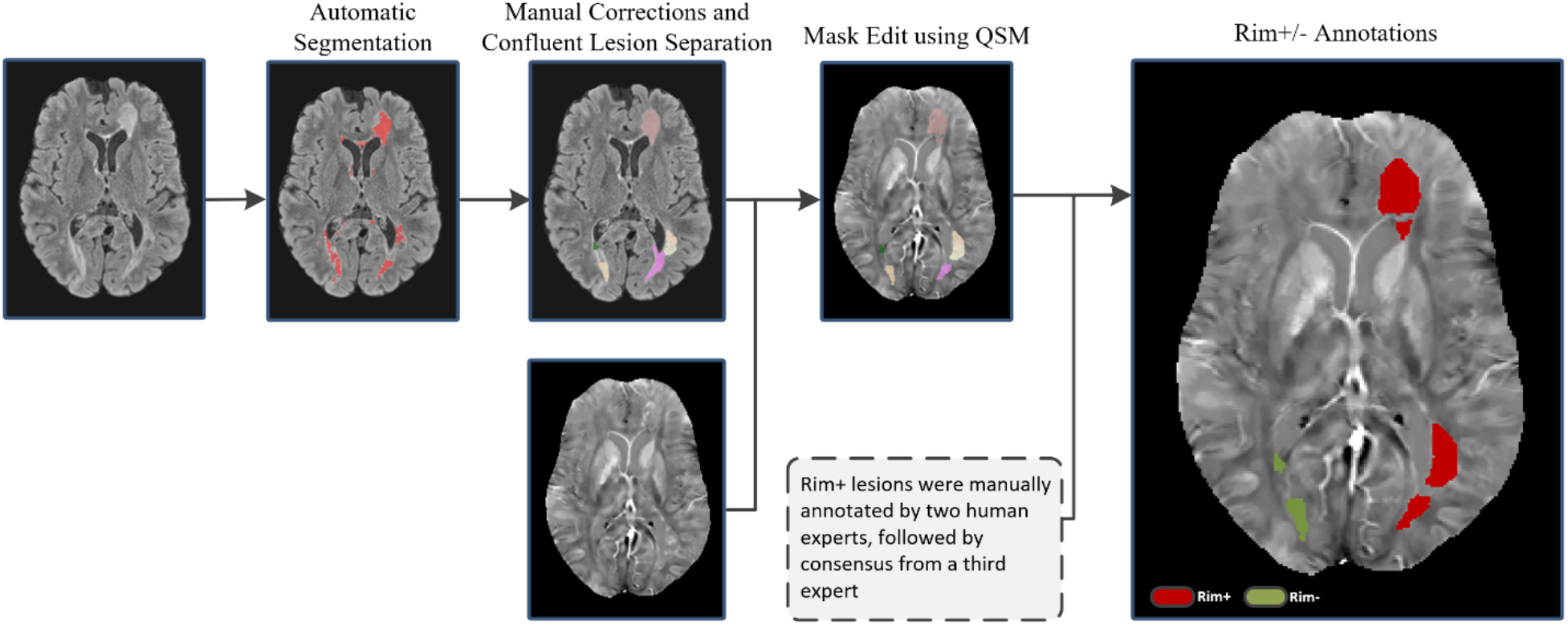
Schematic of the rim+ and rim-lesion annotation process. First, we use LST [34] to obtain an initial lesion segmentation mask. Second, a human expert performs manual correction and confluent lesion separation, followed by mask edits based on QSM. Third, rim+ lesions are manually annotated by two human experts, followed by consensus from a third expert.

Rim+ and rim-lesions were manually annotated by two human experts, who reviewed each of the 4,163 T2-FLAIR lesions for rim status on the QSM. For lesions with disagreement, a final consensus was obtained from a third human expert. After the rim lesion annotation, 177 lesions were identified as rim+ lesions and 3,986 lesions were identified as rim-lesions. An overview of the annotation process is shown in Fig. 2. and the distribution of rim+ lesions per patient is shown in Fig. 3.

**Fig. 3:**
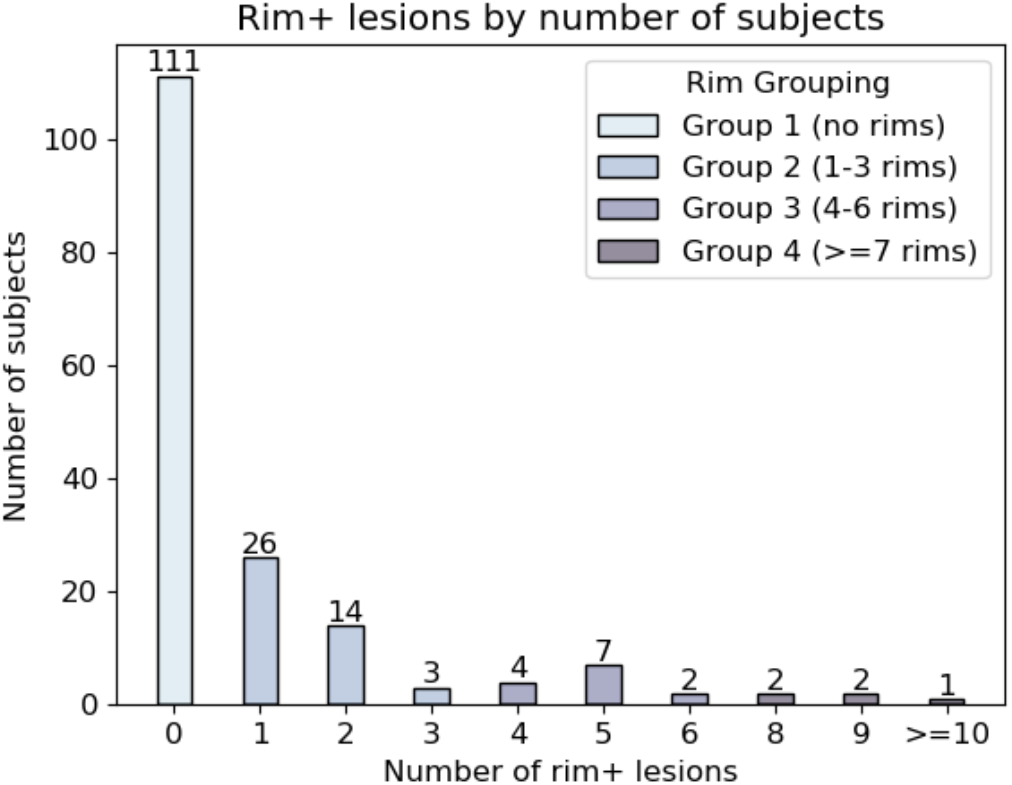
Distribution of the number of paramagnetic rim lesions (rim+ lesions) per subject in the Weill Cornell dataset. The plot is colored by the groups used for the stratified five-fold cross validation.

### 2.2 Methodology

#### 2.2.1 Network architecture

QSMRim-Net is a two-branch network consisting of four parts: a convolutional network for image feature extraction, a fully connected network for radiomic feature extraction, a SMOTE-based oversampling network for synthesizing rim+ features in the latent feature space, and a final classifier that outputs the probability that a lesion is rim+ (see Fig. 4). For image feature extraction, we use a deep residual network (ResNet) [28] with 18 layers as our backbone network. We modified the convolutional kernels from 2D to 3D, used two input channels to accommodate the QSM and T2-FLAIR images, and used two categories (rim+ and rim-) for the last linear layer. For radiomic feature extraction, radiomic features [35, 36] were calculated on the QSM (described in detail in the section below). The multi-layer perceptron (MLP) for radiomic feature extraction consists of two fully connected layers. The first layer is a linear layer followed by a onedimensional batch normalization [37], a Swish activation function [38], and a dropout layer. The second layer has the same structure as the first layer, except that it does not include a dropout layer. To fuse the convolutional and radiomic features, we performed vector concatenation for feature vectors from both the output of the residual network and the MLP and processed the new feature vector with another fully connected layer (see Figs. 4 and 5). To alleviate the data class imbalance issue, we further applied the DeepSMOTE network (described in detail in the section 2.2.3) to oversample these latent features of rim+ lesions during the training phase.

**Fig. 4:**
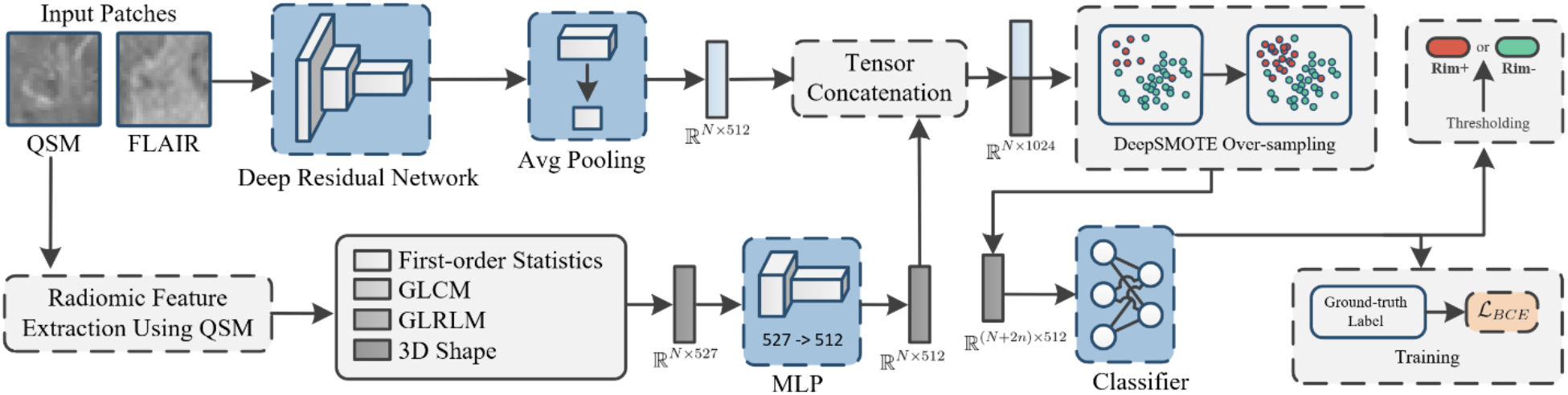
Schematic of the proposed QSMRim-Net for paramagnetic rim lesion identification. (Top Row) The deep residual network takes in both QSM and FLAIR images to extract convolutional features. (Bottom Row) The QSM image and the lesion mask are used to extract radiomic features, followed by feature extraction of an MLP. A tensor concatenation operation is performed to fuse convolutional and radiomic features, and a DeepSMOTE layer is used to perform synthetic minority feature over-sampling during the training phase.

**Fig. 5:**
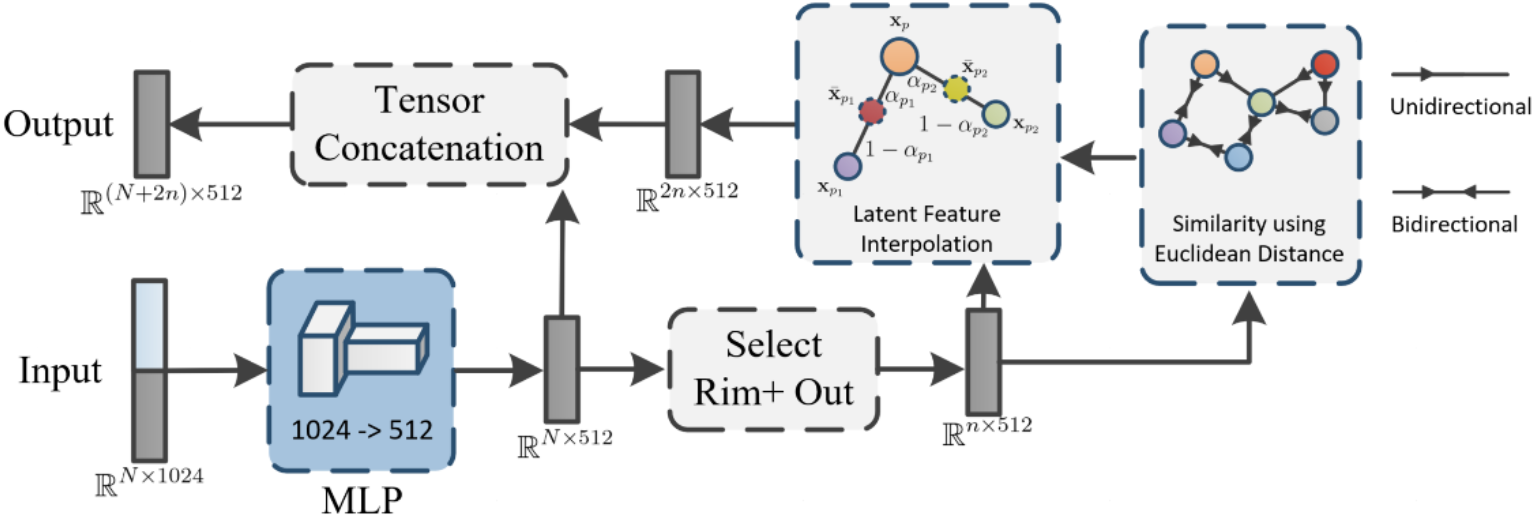
Schematic of the DeepSMOTE network layer. *N* is the number of samples in a training mini-batch, and *n* is the number of rim+ samples in the mini-batch. The input features go through an MLP for feature transformation, followed by selecting rim+ samples from the mini-batch. The transformed rim+ features are used to generate a Euclidean distance-based similarity followed by latent feature interpolation. The original features and the oversampled features are concatenated, resulting in a total of *N* + 2*n* samples in the output of DeepSMOTE.

#### 2.2.2 Radiomic feature analysis

Radiomic features have been shown to be effective in many applications of medical image analysis [40–43]. For QSMRim-Net, radiomic features were calculated over each lesion using the pyradiomics package [44]. Specifically, we calculated four different types of radiomic features on the input image: 1) first-order statistics such as harmonic mean and geometric mean, 2) gray-level co-occurrence matrix (GLCM) statistics such as interquartile range and energy sum, 3) gray-level run-length matrix (GLRLM) statistics such as run-percentage, and 4) geometric-based parameters such as ratio of lesion surface to volume. In addition, Coiflet wavelet filters were applied to yield the 8 decompositions of the input image and radiomic features were calculated on the wavelet images. Wavelet filters were implemented with the PyWavelet package [45]. In total, 527 radiomic features were calculated over each lesion on the QSM for our model.

#### 2.2.3 SMOTE-based oversampling network

Rim+ lesions are rare, with a prevalence of 4.25% in our dataset. This poses a great challenge to training any learning model. To overcome this challenge, we propose DeepSMOTE, a novel oversampling network that leverages the latent features extracted from deep neural network. Intuitively, DeepSMOTE can be thought of as finding the two nearest neighbors of each rim+ lesion and taking a linear combination of the lesion’s features with each of the neighbors’ features to produce synthetic observations. DeepSMOTE consists of a multi-layer perceptron (MLP) for feature transformation followed by the synthetic sample generation. The MLP is designed in a similar style as the network branch for radiomic feature extraction, where there are two consecutive fully connected networks, each having a linear layer, a 1-dimensional batch normalization, and a Swish activation function. The MLP is used to fuse features from the two network branches and reduces the feature dimension from 1024 to 512 for efficient computation (Figs. 4 and 5). Next, the two nearest neighbors for each rim+ lesion in the mini-batch are determined using Euclidean distance. To train QSMrim-Net on a modern GPU, a small portion (mini-batch) of size *N* (64 samples in a mini-batch in our implementation) from the entire training dataset was randomly sampled during one forward-backward pass. Suppose there are *n* rim+ lesions in a particular mini-batch. Let *x_p_* (*p* ∈ {1,2,3… *n*}) be the latent feature vector of a rim+ lesion in the mini-batch. Let *x*_*P*1_ and *x*_*p*2_ be the two nearest rim+ lesions to this lesion in the mini-batch, with respect to Euclidean distance. We generate two synthetic samples:

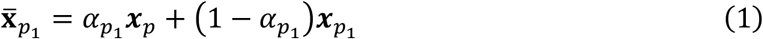

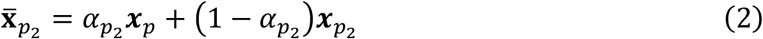

where *α*_*p*1_ and *α*_*p*2_ are randomly generated numbers in (0,1). The result, 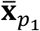 and 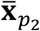, are linear combinations of the rim+ lesion and its nearest neighbors. This results in *N* + 2*n* observations on which to train the network, the *N* samples in the mini-batch and the and the 2*n* synthetic samples. DeepSMOTE differs from the original SMOTE algorithm, which oversamples the minority class by the reciprocal of the percentage of the minority class present in the dataset. DeepSMOTE instead samples 2*n* synthetic samples for each mini-batch during training, as each forwardbackward pass of the deep neural network is done in a mini-batch of the entire dataset and oversampling too many rim+ lesions in a single mini-batch may result in overfitting of the network.

### 2.3 Training details

We applied a stratified five-fold cross-validation procedure to train and validate the performance of QSMRim-Net and the other methods. The stratified procedure was performed to balance the number of rim+ lesions in each of the five folds. As seen in Fig. 3, we first grouped subjects into four groups, where the first group contained subjects with no rim+ lesion, the second subjects with 1-3 rim+ lesions, the third subjects with 4-6 rim+ lesions, and the fourth subjects with more than 6 rim+ lesions. The data was then randomly split into the five folds within each of these groups. All experiments were conducted with this stratified five-fold cross validation setting.

Input images were cropped into image patches with a fixed size 32×32×16 voxels, followed by a masking out of non-lesion area. The largest rim+ lesion had a size of 15×20×5 voxels and only 7 rim- lesions were above the size of 32×32×16. As many lesions are confluent and may overlap in the patch, we only left the target lesion area in the patch. We then performed data augmentation to improve the performance of the model, providing better model generalizability. For augmentation in the training set, lesions were moved to align their center of mass to the geometric center of the image patch. We then used random flipping, random affine transformations, and random blurring to augment our data. Flipping was performed on an orthogonal direction randomly chosen from the axial, coronal, or sagittal direction. Affine transformations were performed with a random scale ranging from 0.95 to 1.05 and a random rotation degree between −5 and 5 degrees. The final transformed patch was obtained after a trilinear interpolation. The blurring was performed using a random-sized Gaussian filter where the kernel radius was determined by 4σ+0.5. The voxel size of our QSM image was 0.75 × 0.75 × 3, thus for the coronal and sagittal direction, we randomly sampled 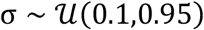, and for the axial direction we randomly sampled 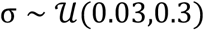.

We implemented our network using the PyTorch library [46] on a computer equipped with a single Nvidia 1080Ti GPU. The Adam algorithm [47], with an initial learning rate of 0.0001 and multi-step learning rate scheduler with milestones at 50%, 70%, and 90% of the total epochs, was used to train the network weights. A mini-batch size of 64 was used for training, and training was stopped after 40 epochs. We used three random seeds to train three models for each fold and the final prediction result was determined by majority voting.

### 2.4 Comparator Methods

Two automated methods have been developed to identify chronic active lesions on phase images [30, 31]. RimNet [30] develops a multi-modal VGGNet [48] to extract rim information from image patches of the phase and T2-FLAIR images. APRL [49] applies a SMOTE and a random forest model to first-order radiomic features derived from individual lesions on the phase images. To evaluate QSMRim-Net, we compared the performance of the proposed algorithm with these two methods. Both methods were originally implemented on the phase, therefore we adapted these methods to a QSM implementation for use with our data. For RimNet, we used the QSM image along with the T2-FLAIR image as the network inputs. For APRL, we used the QSM image to extract the first order radiomic features as done in the original implementation. We applied SMOTE as done in the original APRL method to oversample the rim+ lesion features by the reciprocal of its percentage present in our dataset. In addition to APRL, we also evaluated a neural network with the radiomic features, which is denoted as APRL (NN). The APRL (NN) uses the same network architecture as the radiomic branch of our QSMRim-Net and uses all 527 radiomic features instead of only the first order radiomic features. In the remainder of the manuscript, we refer to the implementation of APRL with the random forest as APRL (RF).

### 2.5 Statistical Evaluation

To evaluate the performance of each method, partial receiver operating characteristic (pROC) curves with false positive rates up to 0.1 and precision-recall (PR) curves of the different validation folds were interpolated using piecewise constant interpolation and averaged to show the overall performance at the lesion-level. For each curve, the area under the curve (AUC) was computed directly from the interpolated and averaged curves. As rim+ lesions are rare and a small subset of the total number of lesions (4.25% of the lesions), allowing a high false positive rate threshold would produce results that are not clinically relevant. We therefore examine the pROC for false positive rates between 0 and 0.1. In addition, to create binary maps of rim+ versus rim-lesions, we thresholded the model probabilities to optimize the F1 score. The F1 score is the harmonic mean of precision and sensitivity, where 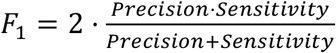. The thresholds for different folds of the cross-validation were chosen separately, and all results were obtained by a concatenation of the results for the five folds. For the lesion-wise analysis, accuracy, F1 score, sensitivity, specificity, and positive predicted value (PPV) were used to characterize the performance of each automated method.

We also assessed performance at the subject-level. We used the F1 score criteria for thresholding and compared the number of predicted rim+ lesions and the expert count number of rim+ lesions for each subject. Pearson’s correlation coefficient was used to measure the correlation between the two values. Mean Squared Error (MSE) was also used to measure the averaged accuracy for the model predicted count.

### 2.6 Ablation study

We conducted an ablation study to evaluate the effects of changing components of the QSMRim-Net network using ResNet as the backbone network. First, we examined two different fusion methods for the image-level features. In RimNet, a multi-modal architecture was used to fuse image-level features. In QSMRim-Net we concatenated images in the channel dimension of the tensor for input into the network. Second, we investigated the effects of incorporating the radiomic feature as a separate network branch. Third, we studied how the DeepSMOTE layer affects the network performance. The following four schemes were evaluated based on the method for fusing the QSM and T2-FLAIR imaging features and whether we adopt radiomic feature or DeepSMOTE layer: 1) images were fused as in QSMRim-Net with no radiomic features, 2) images were fused as in RimNet with no radiomic features, 3) images were fused as in QSMRim-Net with radiomic features, 4) images were fused as in QSMRim-Net with radiomic feature and DeepSMOTE network was adopted.

## 3. RESULTS

### 3.1 Lesion-wise analysis

Table 2 shows the lesion-wise performance metrics of the proposed QSMRim-Net and the other methods, using the F1-score as a threshold. QSMRim-Net outperformed the competitors in all metrics used for evaluation. With a slightly higher overall accuracy and specificity with other methods, QSMRim-Net resulted in a 9.8% and 23.3% improvement in F1 score, 3.5% and 14.3% improvement in sensitivity and 16.8% and 33.1% improvement in PPV compared to Rim-Net [30] and APRL (RF) [46], respectively.

Fig. 6 shows the pROC curves and the PR curves for the different methods. The proposed QSMRim-Net obtained 4.68% and 21.01% higher pROC AUC (0.760) than Rim-Net (0.726) and APRL (RF) (0.628), meaning that for more clinically relevant false positive rates of less than 0.1, QSMRim-Net has higher performance than the other methods. The proposed QSMRim-Net out-performed both Rim-Net and APRL (RF) by 9.8% and 60.0% respectively in PR AUC, indicating the effectiveness of the proposed DeepSMOTE network and fusion of information from both convolutional image and radiomic features. Interestingly, APRL (NN) that uses our neural network architecture outperformed APRL (RF) significantly in both pROC AUC and PR AUC, indicating the potential for the neural network to exploit high-dimensional non-linear relationships from the radiomic features.

**Fig. 6:**
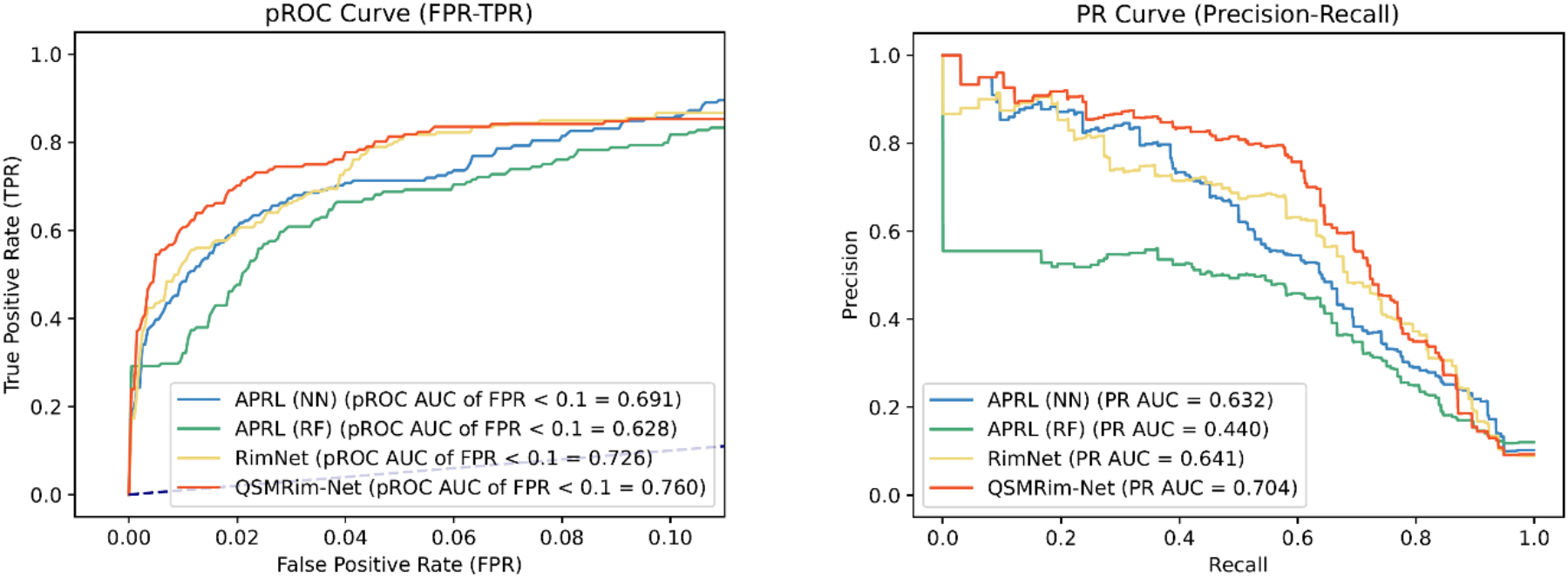
The partial receiver operating characteristic (pROC) curve and precision-recall (PR) curves for the proposed (QSMRim-Net) and comparator methods. AUC denotes the area under the curve. We use clinically relevant false positive rates of less than 0.1 to compute the pROC AUC, in order to account for the rare nature of rim+ lesions. Our QSMRim-Net algorithm outperformed all other algorithms on pROC AUC (FPR < 0.1) and PR AUC.

### 3.2 Subject-wise analysis

We calculated the predicted count of rim+ lesions from each of the models and compared this to consensus expert count for each subject. The consensus expert count of rim+ lesions ranged from 0 to 17 among the 172 subjects, with a median of 2 rim+ lesions among subjects with at least one rim (IQR 1–4). The predicted count of rim+ lesions from QSMRim-Net ranged from 0 to 14, with a median of 1 rim+ lesion among the subjects with at least one rim (IQR 1–4).

The Pearson’s correlation between the predicted count and the gold standard count was 0.89 (95% CI: 0.86, 0.92). Fig. 7 shows the scatterplot for the predicted count versus the gold standard count, along with the identity line. The Pearson’s correlations for the other methods were found to be lower than QSMRim-Net: 0.88 (95% CI: 0.85. 0.91) for APRL (NN), 0.77 (95% CI: 0.70, 0.82) for APRL (RF), and 0.75 (95% CI: 0.67, 0.81) for Rim-Net. The MSE for the predicted count of the QSMRim-Net was 0.98. The MSE for the other methods were found to be higher: 1.02 for APRL (NN), 2.26 for APRL (RF), and 2.47 for Rim-Net.

**Fig. 7:**
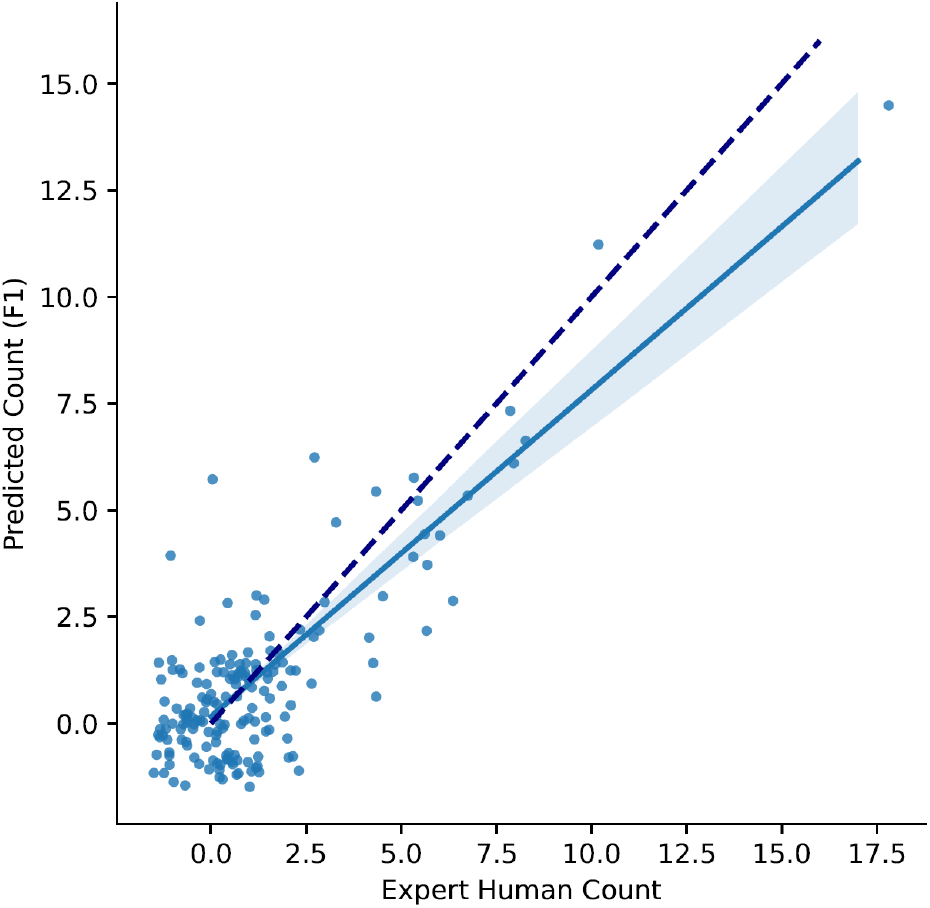
The predicted count of rim+ lesions from QSMRim-Net versus the expert human count (*ρ* = 0.89 (95% *CI*: 0.86,0.92), *MSE* = 0.98). Points in the plot have been jittered for better visualization. The linear regression line for the predicted count versus the gold standard count with 95% CI is also shown (solid blue) along with the identity line (dashed blue).

### 3.3 Ablation study

Table 3 shows the results from the ablation study. For image fusion, the fusion technique in QSMRim-Net outperformed the fusion technique for RimNet. For radiomic feature fusion, we can see that the network with radiomic feature fusion performs better than the networks without radiomic feature fusion. The network with Deep SMOTE oversampling outperforms the other variants in all metrics.

## 4. DISCUSSION

In this paper, we propose QSMRim-Net, a deep convolutional neural network for identifying rim+ lesions on QSM MRI. This is the first algorithm developed for rim+ lesion identification on QSM and the first study to introduce an end-to-end two-branch network enabled with the DeepSMOTE oversampling technique that can effectively fuse both convolutional image and radiomic features.

Our QSMRim-Net achieved better performance than two previously developed methods when applied to the QSM, APRL (RF) [49] and Rim-Net [30]. The increase in performance can be attributed to our carefully designed convolutional neural network architecture (Fig. 4). Rim-Net used VGG-Net [48] and APRL (RF) used a random forest. Our QSMRim-Net adopted a ResNet [28] architecture that uses identity shortcut connections to prevent gradient vanishing, which reduces computational complexity and allows for the training of deeper networks than VGG-Net. We also observed that a neural network with MLP (APRL (NN)) achieved better performance than a random forest model (APRL (RF)) on radiomic features. This shows that a properly designed neural network can extract discriminative information from highly non-linear radiomic feature data. Another contributor to QSMRim-Net’s performance is that it effectively fuses the complementary information from the convolutional image and radiomic features. We also showed in the ablation study that the neural network architecture design choices for fusing features from different sources is important for improving rim+ lesion identification performance.

In addition to the deep neural network model, our result may also benefit from the utilization of QSM. Compared to the phase images used in the original implementation of Rim-Net and APRL (RF), QSM can measure the underlying tissue apparent magnetic susceptibility, enabling the quantification of specific biomarkers, such as iron, that are independent of imaging parameters. Rim+ lesions are characterized by a paramagnetic rim with iron deposited at the edge of the lesion. QSM is sensitive to such magnetic susceptibility changes and provides consistent measurements of the susceptibility value of the rim across patients and scanners, which is beneficial for a machine learning model such as deep neural network to learn patterns of rim+ and rim-lesions. While our QSMRim-Net is inspired by Rim-Net and APRL (RF), we found that implementing these two methods on our dataset using QSM resulted in 10.2% and 25.0% improvement of PPV, 7.8% and 26.5% reduction in sensitivity respectively, compared to its original implementation of Rim-Net and APRL (RF) on phase images.

The high imbalance of rim+ and rim-lesions is a challenge for machine learning models. We found that APRL (RF) with SMOTE oversampling outperforms its counterpart without SMOTE by 2.9% in F1 score, indicating the importance of oversampling of the minority class. While it is feasible to synthesize radiomic features by linear interpolation using SMOTE, it is not possible to synthesize meaningful images by pixel-level linear interpolation. It has been shown empirically that deep neural networks can linearize the manifold of images into Euclidean subspace [50–52], enabling the possibility of linear interpolation using latent features from deep layers from the network. Inspired by the SMOTE results from APRL (RF) and the deep feature interpolation, we propose DeepSMOTE network to alleviate the data imbalance issue, and the results in Table 3 shows the effectiveness of applying DeepSMOTE for data oversampling.

For the QSMRim-Net performance results, we thresholded the output probabilities from the algorithm using the optimized F1 score, resulting in a sensitivity of 0.678 for detecting rim+ lesions. However, in a research scenario or in clinical practice, missing any rim+ lesions may not be acceptable. Thus, to demonstrate the performance of QSMRim-Net in these settings, we also applied a high sensitivity threshold, using the largest sensitivity below 0.95. In practice, experts can use this high sensitivity threshold to reduce the number of lesions that need to be manually reviewed for rim+ status. QSMRim-Net performed with a false positive number of 538 lesions, a reduction of 14.2% and 19.1% compared to Rim-Net and APRL (RF). With QSMRim-Net, in this dataset, only 715 lesions would need to be reviewed by an expert instead of all 4163 lesions, saving 82.8% of review time.

We also obtained results for all rim+ lesion identification algorithms on a patient-level, showing that QSMRim-NET outperformed the other methods. In a previous study, as the overall total lesion burden increased, patients with at least one rim+ lesion on QSM performed worse on both physical disability and cognitive assessments [17]. Having four or more chronic active lesions on phase imaging has also been shown to correlate with disability [16]. In addition, these lesions have been used diagnostically to differentiate patients with MS from other neurological conditions [53]. If rim+ lesions are to be used prognostically or diagnostically, then the patient-level results may be more important than identifying individual rim+ lesions for clinical translation.

To further understand the limitations of the QSMRim-Net algorithm’s performance, we also examined the false positive and false negative results. The false positive and negative results tended to be lesions that the two human experts did not agree upon. Using the F1 score threshold, 40.9% of the false positive lesions and 35.1% of false negatives lesions were lesions with human expert disagreement. This contrasts with 22.5% of the true positives and 2.4% of the true negatives.

Visual examination (Fig. 8) of the lesions showed that veins were challenging for the algorithm, resulting in false positives. On QSM the vein is hyperintense. In cases where the vein formed a rim-like shape in a rim-lesion, this often resulted in a false positive (Fig. 8B). When a rim+ lesion was found heterogeneously hyperintense, this often resulted in a false negative (Fig. 8C). Rim-lesions with a higher intensity value on the QSM tended to cause false positives, while rim+ lesions with a lower intensity value tended to cause false negatives. Our future work involves further understanding of these patterns to reduce FPs and FNs from the algorithm in order to improve research and clinical translation.

**Fig. 8:**
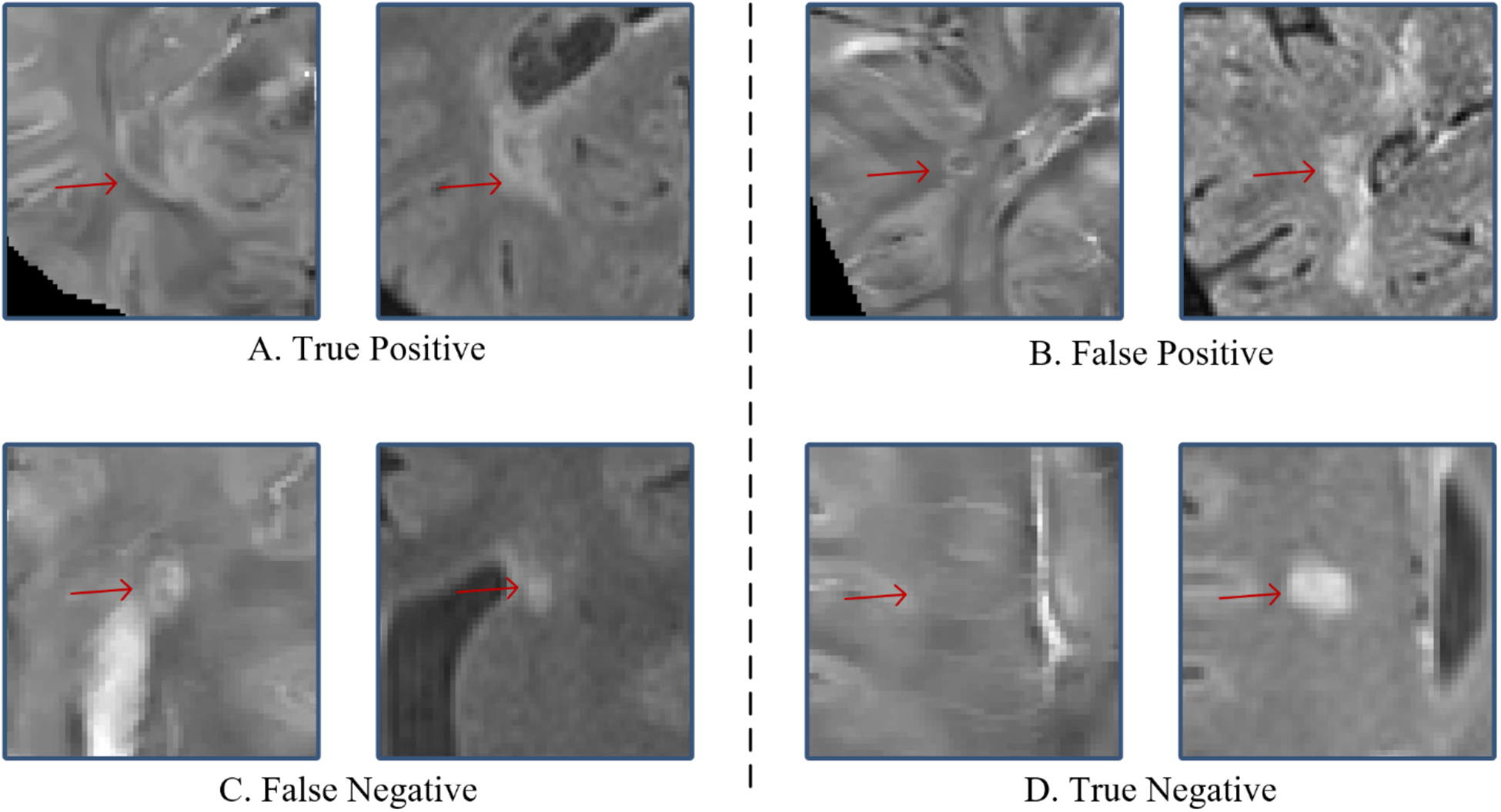
Visual examples of a true positive, a false positive, a false negative and a true negative produced by QSMRim-Net. The QSM is shown on the left and the T2-FLAIR on the right. The lesion of interest indicated with a red arrow. (A) A rim+ lesion that is correctly identified. (B) A rim-lesion with a vein forming a rim-like shape that was falsely identified as rim+ by QSMRim-Net. (C) A rim+ lesion with that was missed by QSMRim-Net. (D) A rim-lesion that is correctly identified.

One limitation of this work is that it relies on manual lesion segmentations that have been edited further on the QSM (Fig. 2). Future work involves pairing QSMRim-Net with an automated T2-FLAIR lesion segmentation algorithm, such as All-Net [21] with geometric loss [54] and attention-based approaches [55] [56], followed by an automated method to separate confluents lesions [57]. We plan to adapt and train the algorithm to work directly on T2-FLAIR lesion segmentations. Another challenge for the algorithm is the rare nature of rim+ lesions. Only 4.25% of the lesions in this study were identified as rim+ lesions, posing a great challenge to learning based methods. We proposed DeepSMOTE for data oversampling to alleviate the data class imbalance, but as future work we plan to develop techniques on imbalance-aware loss functions, such as geometric loss [54]. A further limitation is inter-rater variability in identifying rim+ lesions. To reduce the impact of this, we had two raters evaluate lesions for rim+ status and any disagreements were adjudicated by a third reviewer. In this work, we used a binary classification of whether a lesion had a rim. As discussed in [49] there are many factors that influence the strength of the rim+ lesion signature on QSM and a more nuanced approach to classify these lesions may be beneficial.

In conclusion, QSMRim-Net is the first deep learning-based method that integrates DeepSMOTE for data oversampling and fuses modern convolutional imaging features with traditional radiomic features to automatically identify rim+ MS lesions on QSM. QSMRim-Net out-performed other state of the art methods on rim+ lesion identification on QSM and has the potential to aid in the clinical translation for the rim+ lesion biomarker.

**Table 1.** Demographics information for the study cohort.

| Demographics |  |
| --- | --- |
| Number of Subjects | 172 |
| Gender (count (%)) |  |
| Female | 124 (72.09%) |
| Male | 48 (27.91%) |
| Disease Subtypes (count (%)) |  |
| RRMS | 159 (92.44%) |
| SPMS | 8 (4.65%) |
| CIS | 5 (2.91%) |
| Disease Durations (mean ± STD) | 10.68 ± 7.37 |
| Age (mean ± STD) | 42.82 ± 10.27 |
| Expanded Disability Status Score (mean ± STD) | 1.38 ± 1.64 |
| Treatment Durations (mean ± STD) | 8.05 ± 5.79 |

**Table 2.** Lesion-wise results of the QSMRim-Net and other methods using a stratified five-fold cross-validation scheme. PPV denotes positive predictive value, TP# denotes the number of true positives, FP# denotes the number of false positives, FN# denotes the number of false negatives, and TN# denotes the number of true negatives. The best performing method for each of metrics is bolded.

| Lesion-wise results | Accuracy | F1 | Sensitivity | Specificity | PPV | TP # | FP # | FN # | TN # |
| --- | --- | --- | --- | --- | --- | --- | --- | --- | --- |
| APRL (NN) | 0.969 | 0.614 | 0.588 | 0.985 | 0.642 | 104 | 58 | 73 | 3928 |
| APRL (RF) | 0.962 | 0.571 | 0.593 | 0.978 | 0.550 | 105 | 86 | 72 | 3900 |
| Rim-Net | 0.969 | 0.641 | 0.655 | 0.983 | 0.627 | 116 | 69 | 61 | 3917 |
| QSMRim-Net | **0.976** | **0.703** | **0.680** | **0.989** | **0.732** | **120** | **44** | **57** | **3942** |

**Table 3.** Ablation study on QSMRim-Net and its variants. PPV denotes positive predictive value, TP# denotes the number of true positives, FP# denotes the number of false positives, FN# denotes the number of false negatives, and TN# denotes the number of true negatives. Image Fusion indicates whether the model performed image-level feature fusion, Radiomic Fusion indicates whether the model performed feature fusion between image and radiomic features, and Deep SMOTE indicates whether the model applied the DeepSMOTE network for rim+ feature oversampling. The best performing method for each of metrics is bolded.

| Image<br>Fusion | Radiomic<br>Fusion | Deep<br>SMOTE | Accuracy | F1 | Sensitivity | Specificity | PPV | TP# | FP# | FN# | TN# |
| --- | --- | --- | --- | --- | --- | --- | --- | --- | --- | --- | --- |
| ✓ | × | × | 0.969 | 0.639 | 0.644 | 0.983 | 0.633 | 114 | 66 | 63 | 3920 |
| × | × | × | 0.971 | 0.645 | 0.610 | 0.987 | 0.684 | 108 | 50 | 69 | 3936 |
| × | ✓ | × | 0.975 | 0.685 | 0.650 | **0.989** | 0.723 | 115 | **44** | 62 | **3942** |
| × | ✓ | ✓ | **0.976** | **0.703** | **0.680** | **0.989** | **0.732** | **120** | **44** | **57** | **3942** |

## Notes

### Competing Interest Statement

The authors have declared no competing interest.

